# Quantification of Mouse Reach Kinematics as a Foundation for Mechanistic Interrogation of Motor Control

**DOI:** 10.1101/2020.04.24.060533

**Authors:** Matthew I. Becker, Dylan Calame, Julia Wrobel, Abigail L. Person

## Abstract

Mice use reaching movements to grasp and manipulate objects in their environment, similar to primates. Thus, many recent studies use mouse reach to uncover neural control mechanisms, but quantification of mouse reach kinematics remains lacking, limiting understanding. Here we implement several analytical frameworks, from basic kinematic relationships to statistical machine learning, to quantify mouse reach kinematics across freely-behaving and head-fixed conditions. Overall, we find that many canonical features of primate reaches are conserved in mice, with some notable differences. Our results highlight the decelerative phase of reach as important in driving successful outcome. Late-phase kinematic adjustments are yoked to mid-flight position and velocity of the limb, allowing dynamic correction of initial variability, with head-fixed reaches being less dependent on position. Furthermore, consecutive reaches exhibit positional error-correction but not hot-handedness, implying opponent regulation of motor variability. Overall, our results establish foundational mouse reach kinematics in the context of neuroscientific investigation.

## Introduction

As a primary mode of interaction with the environment, reaching movements are a fundamental motor behavior across species. Because reaches are rapid and require coordination across multiple joints, they implicate neural processes such as internal models and predictive state estimates, ideas that have been highly influential in neuroscience (Reza Shadmehr et al., 2010; Reza Shadmehr & Krakauer, 2008; Wolpert et al., 1995). Moreover, restoring limb function in humans suffering from stroke or neurological injury is an important topic in rehabilitative medicine, driving research in motor control in service of brain computer interface technology (Pandarinath, Ames, et al., 2018; Pandarinath, O’Shea, et al., 2018; Schroeder & Chestek, 2016; Stavisky et al., 2017). Much of the scientific literature on reaching movements has been performed in primates, giving rise to many prominent motor control theories (Fitts, 1954; Harris & Wolpert, 1998b; R Shadmehr & Mussa-Ivaldi, 1994). From motor control models, such as minimum jerk and optimal control, to neural system mechanisms, such as preparatory activity and dynamical systems, reaching movements served as an essential model system for advancing neuroscientific investigation (Flash & Hogan, 1985; Shenoy et al., 2013; Tanji & Evarts, 1976; Todorov & Jordan, 2002). While many of these computational frameworks derive from human psychophysical work, identifying neural correlates and mechanisms in non-human primates remains a core focus of neurophysiological investigations.

Reaching movements are evolutionarily conserved, however, opening up the possibility of interrogating these movements using the advanced molecular, cell-type specific, and neural circuit interrogation methods readily available in mice (Iwaniuk & Whishaw, 2000). Whishaw and colleagues developed a self-initiated forelimb reach-to-grasp task for rodents, in which freely-behaving mice retrieve small pellets of food from a shelf, accessible by the limb through a small opening in a behavioral arena (Whishaw, 1996; Whishaw & Pellis, 1990). Over the last decade, the study of mouse reach has grown considerably, and the task has been adapted in several ways, including head-fixed conditions, use of levers or joysticks, conditioned stimuli for reach initiation, and grasping of various objects including water (Azim, Fink, et al., 2014; Bollu, Whitehead, Prasad, et al., 2019; Esposito et al., 2014; Galiñanes et al., 2018a; Guo et al., 2015; Mathis et al., 2017; Miri et al., 2017; Peters et al., 2014; Yttri & Dudman, 2016). Despite the recent prominence of mouse forelimb movement studies, a relative dearth of quantitative analyses of mouse reach kinematics precludes mechanistic interrogation of the well-developed motor control principles discussed above. Specifically, in both human and nonhuman primates, careful quantitative descriptions of reaching movements have provided a crucial foundation for understanding neurophysiological control mechanisms.

Ethologically, the purpose of reaching movements is to grasp and manipulate objects, goals which constrains how reaching movements progress through time and space (i.e. reach kinematics). In primates, repeated observations of common kinematic features, such as the bell-shaped velocity profile or the intrinsic relationship between speed, distance, and duration, have provided insight into underlying neural mechanisms of reach production (Flash & Hogan, 1985; Harris & Wolpert, 1998b, 2006; Jeannerod, 1984; Meyer et al., 1988; Morasso, 1981). Because neural circuits underlying reach behavior exert explicit control over task-relevant variables (Domkin et al., 2005; Osu et al., 2015; Scholz & Schöner, 1999), which can vary across conditions as goals and constraints change, there is a need to identify and quantify kinematic correlational structure that characterizes mouse reach behavior. Of particular importance for reach-to-grasp success is endpoint positioning. Indeed, the earliest conceptions of reach behavior hypothesized a two-part model of control, involving an initial, centrally-driven ‘impulse’ phase designed to bring the limb near the target, and a second, feedback-based ‘homing’ phase designed to impart final corrections that ensure accurate endpoints (Elliott et al., 2001; Woodworth, 1899). Powerful modern motor control frameworks now include explicit models that minimize movement cost, incorporate feedback and noise, and can be flexibly altered by task goals, providing an improved ability to explain commonly-observed late-phase alterations in reach kinematics (Diedrichsen et al., 2010; Flash & Hogan, 1985; Meyer et al., 1988; Pruszynski & Scott, 2012; Todorov & Jordan, 2002). In mice, recent work found an adaptive scaling of cerebellar limb control signals during the decelerative phase of reach (Becker & Person, 2019), potentially providing a neurophysiological mechanism for late-phase kinematic corrections observed in humans. Nevertheless, identifying task-relevant control parameters as well as the basic correlational structure of mouse reach kinematics remains in its infancy, motivating the current study.

Here we define foundational kinematic metrics that characterize behaviorally salient aspects of mouse reach behavior. To do so, we track paw kinematics as mice reach to grasp and retrieve small pellets of food from a pedestal, either in a freely-behaving or head-fixed context. Taking inspiration from the primate reach literature, we find that mouse reach kinematics obey well-known relationships, including bell-shaped speed profiles, the speed-accuracy tradeoff, and a tripartite relationship between speed, distance, and duration. We next confirm that endpoint accuracy dictates reach outcome and use a statistical machine learning algorithm to identify kinematic signatures of successful reaches. Reach kinematics proximal to endpoint were the most important for predicting reach outcome, directly implicating late-phase adjustments to initial kinematic variability as critical to performance. Focusing on deceleration from peak velocity to endpoint, we characterize an adaptive scaling of mean deceleration based on both position and velocity parameters, with a stronger dependence on position in the freely-behaving condition. Finally, trial-over-trial analyses suggest errorcorrection mechanisms engaged on consecutive reach attempts, but show a lack of hot-handedness across sessions, indicating a competitive balance between processes that drive motor variability versus maintain precision. We discuss how continued investigation of the neural and biomechanical underpinnings of these reach behavioral features will bolster reach as a canonical model behavior for systems neuroscience, with implications for this behavior across evolutionarily diverse species.

## Results

### Kinematic influences on reach outcome

Mouse reach is an increasingly prevalent behavioral paradigm in the study of motor control. Even so, quantitative descriptions of mouse reach kinematics remain lacking. Defining mouse reach kinematic control parameters, including intrinsic kinematic relationships and their effects on reach accuracy, is a prerequisite for interrogating the neural mechanisms underlying reach behavior in this species. To track mouse reach kinematics, we leveraged machine vision motion capture technology that accurately measures the 3dimensional position of the paw with high spatiotemporal precision (Figure 1A-C). Freely-behaving mice were trained on a self-initiated reach task to retrieve small pellets of food from a pedestal (see Methods; N = 26 animals, 550 recording sessions). We segmented the tracking data into individual reaches, defined as movements of the limb outside of the behavioral arena, and focused on the ‘outreach’ phase of the movement, from reach start until reach endpoint (Figure 1D,E; mean n = 612 reaches, range 205 – 1653; total 15902 reaches). For analysis, we collated reaches on a per-animal basis and extracted average kinematic parameters of interest (Figure 1E; reach start to endpoint; see Methods), including endpoint location, distance travelled, peak velocity, duration, and mean deceleration.

**Figure 1.**
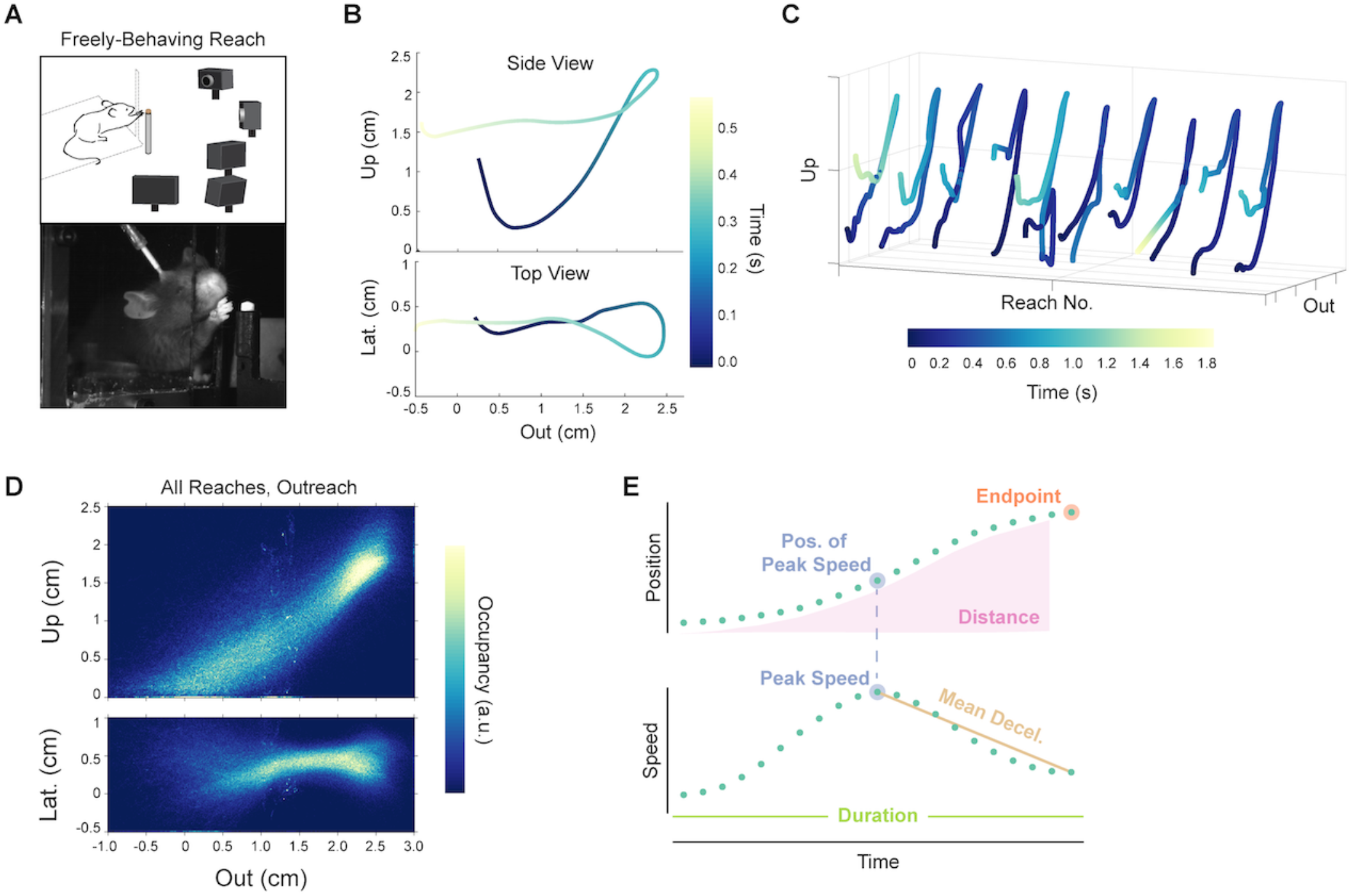
Tracking and quantifying mouse reach behavior. A. Behavioral setup. Top, schematic of freely-behaving reach tracking behavioral arena. Five cameras track the position a small reflective marker affixed to the paw as the animal performs reaching movements to retrieve small pellets of food from a pedestal. Bottom, image of a mouse performing a reaching movement. B. Example mouse reach trajectory (reproduced from Becker & Person, 2019). Position is plotted from a side view (top) and top view (bottom) with time indicated in color. C. Several example reach trajectories. D. Heatmap of positional occupancy of the tracking marker over all 15902 reaching movements, outreach only. Position is plotted from a side view (top) and a top view (bottom). E. Schematic of kinematic parameters of interest.

We began by examining reach endpoint, a kinematic feature that is clearly central to behavioral performance. To obtain the object, reaches must be accurately targeted to the correct final location to enable grasping. We define an endpoint as a direction reversal along the major axis of the reach, which in our task is outward from the animal to the pellet. In order to examine whether endpoint is a key kinematic control variable, we conducted two tests. First, we asked whether mean endpoint position was different on successful versus unsuccessful reaches, which would represent a change in reach accuracy. Second, we asked whether unsuccessful reach endpoints were more variable, which would represent a change in reach endpoint precision. We found that more than half of animals tested displayed differences in reach endpoint location in the outward direction on successful versus unsuccessful attempts (16/26 animals, Wilcoxon rank sum, p < 0.05). Of these animals, most (11/16) showed hypometric mean endpoints on failures compared to successes (Figure 2A), which has been observed in human reaching as well (Chua & Elliott, 1993; Julie Messier & Kalaska, 1997; K. E. Novak et al., 2003). Across all animals, mean endpoint position on failures was hypometric compared to success, although the effect size was marginal (Figure 2B, left; Position from origin; Success: 2.53 cm; Failure: 2.52 cm; Wilcoxon signed-rank, p = 0.046). Though the average endpoint locations of successes and failures were similar, failed reaches displayed significantly higher endpoint variability (25/26 animals, Levene’s Test, p < 0.05). Standard deviation of reach endpoint was much higher for unsuccessful reaches than successful reaches across the population of animals (Figure 2B, right; standard deviation; Success: 0.13 cm; Failure: 0.19 cm; Wilcoxon signed-rank, p < 0.0001). Overall, unsuccessful reaches skewed toward undershooting the target and were more variable in endpoint positioning, confirming endpoint position and variability as key parameters underlying skilled reach.

**Figure 2.**
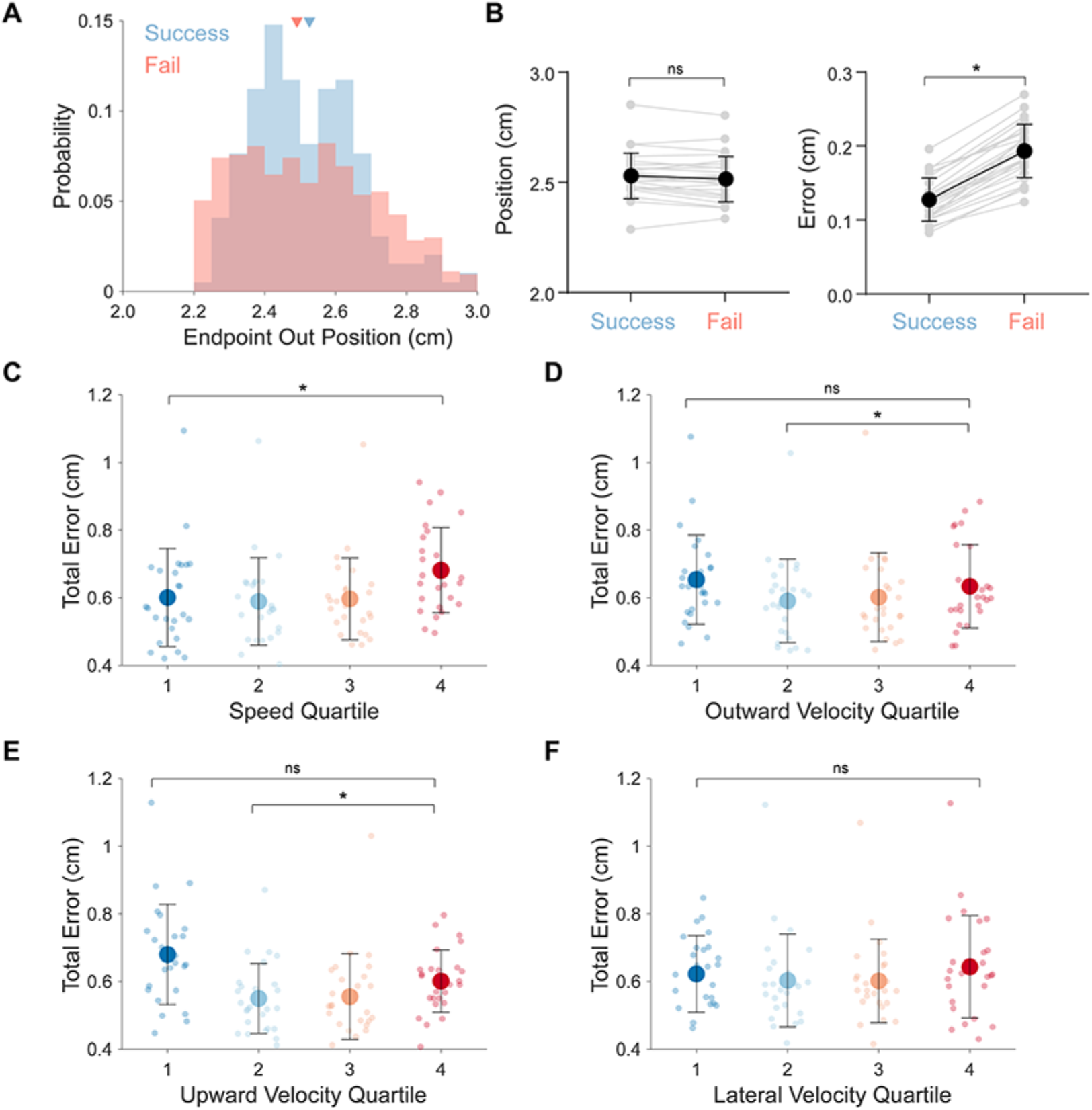
Mouse reach endpoint statistics related to reach outcome and speed. A. Distributions of endpoints in the outward dimension on successful (blue) and failed (red) reaches from an example animal. Group means are indicated by arrowheads. B. Individual animal (gray) and grand mean (black) average endpoint position (left) and standard deviation (right) in the outward dimension for successful and failed reaches. C. Individual animal (small points) and grand mean (large points) total error of endpoints for reaches grouped into quartiles according to peak speed. Reaches were grouped on a per-animal basis. Total error is defined as the sum of the standard deviations calculated individually for each positional dimension (outward, upward, lateral). D-F. Individual animal (small points) and grand mean (large points) total error of endpoints for reaches grouped into quartiles according to peak outward velocity (D), peak upward velocity (E), and peak lateral velocity (F).

Next, we asked how reach speed affects reach accuracy. From a behavioral perspective, faster reaches result in faster retrieval of the object. However, in primates, increased reach speed is associated with decreased accuracy, known as the speed-accuracy tradeoff, a relationship hypothesized to result from signal-dependent noise in the motor system (Fitts, 1954; Harris & Wolpert, 1998a; Meyer et al., 1988; K. E. Novak et al., 2003; Woodworth, 1899). To test for a speed-accuracy tradeoff in mice, we separated reaches into quartiles based on peak speed, and analyzed the cumulative three-dimensional error of endpoints. Reaches in the top speed quartile had higher endpoint error than reaches in the bottom speed quartile, although the effect size was modest (Figure 2C; Top: 0.60 +/− 0.15 cm; Bottom: 0.68 +/− 0.13 cm; Wilcoxon signed-rank p = 0.02). To inspect this relationship further, we decomposed speed into its three Euclidean velocity vectors. In contrast to speed, velocity did not scale monotonically with endpoint error: reaches in the top and bottom quartiles of peak outward velocity showed no difference in overall endpoint error (Figure 2D; Top: 0.63 +/− 0.12 cm; Bottom: 0.65 +/− 0.13 cm; Wilcoxon signed-rank, p = 0.35). Instead, reaches that had outward velocities near average (25th to 75th percentiles) showed decreased endpoint error, revealing that extremes of peak outward velocity (e.g. slow or fast reaches) are associated with decreased accuracy (Figure 2D; Middle-Low: 0.59 +/− 0.12 cm; Middle-Low vs. Top, Wilcoxon signed-rank, p = 0.046). Grouping by upward velocity showed a similar trend, with significantly higher endpoint error in reaches in the lowest and highest quartiles (Figure 2E; Middle-Low vs. Top, p = 0.03). In contrast, organizing reaches by peak lateral velocity resulted in no significant differences in endpoint error across quartiles (Figure 2F; all pairs, Wilcoxon signed-rank, p > 0.05), possibly due to minimal lateral movement in our task. These results confirm that accuracy decreases with overall reach speed, at least for the fastest reaches, with outward and upward velocity being most contributory to differences in endpoint accuracy. The data also point to reach vigor being associated with effective control, as reaches that are too slow or too fast are less accurate, suggesting a ‘sweet spot’ for well controlled reaching. Together, we identified both endpoint location and reach speed as control parameters that influence reach outcome.

### Machine learning for unbiased identification of kinematic contributors to reach outcome

Endpoint location and reach speed have clear and well-demonstrated effects on reach outcome. However, other kinematic variables may also influence the likelihood of success or failure, potentially in non-intuitive ways. For example, the same kinematic variable may have different influences on outcome over time as the reach progresses. We therefore developed a statistical machine learning model to analyze the importance of reach kinematic parameters (position, velocity, and acceleration in three dimensions) in generating successful or unsuccessful outcomes. Our approach modeled reach trajectories in four time segments from start to endpoint, which served as inputs into a random forest prediction algorithm (Figure 3A,B). This allowed us to analyze distinct temporal components of kinematic variables in a highly nonlinear fashion, capturing complex interactions both among kinematic variables and over time. A decision tree (Figure 3B, left) split kinematic variables at certain values to determine a decision pathway that best predicted success or failure for each reach. Hundreds of these trees were generated using random samples of the training data to create a forest (Figure 3B, right), and predictions were averaged across trees in the forest. This random forest model provides several outputs, including overall predictive accuracy, a quantitative assessment of the importance of each kinematic variable segment to the prediction, and a probability of success for each reach.

**Figure 3.**
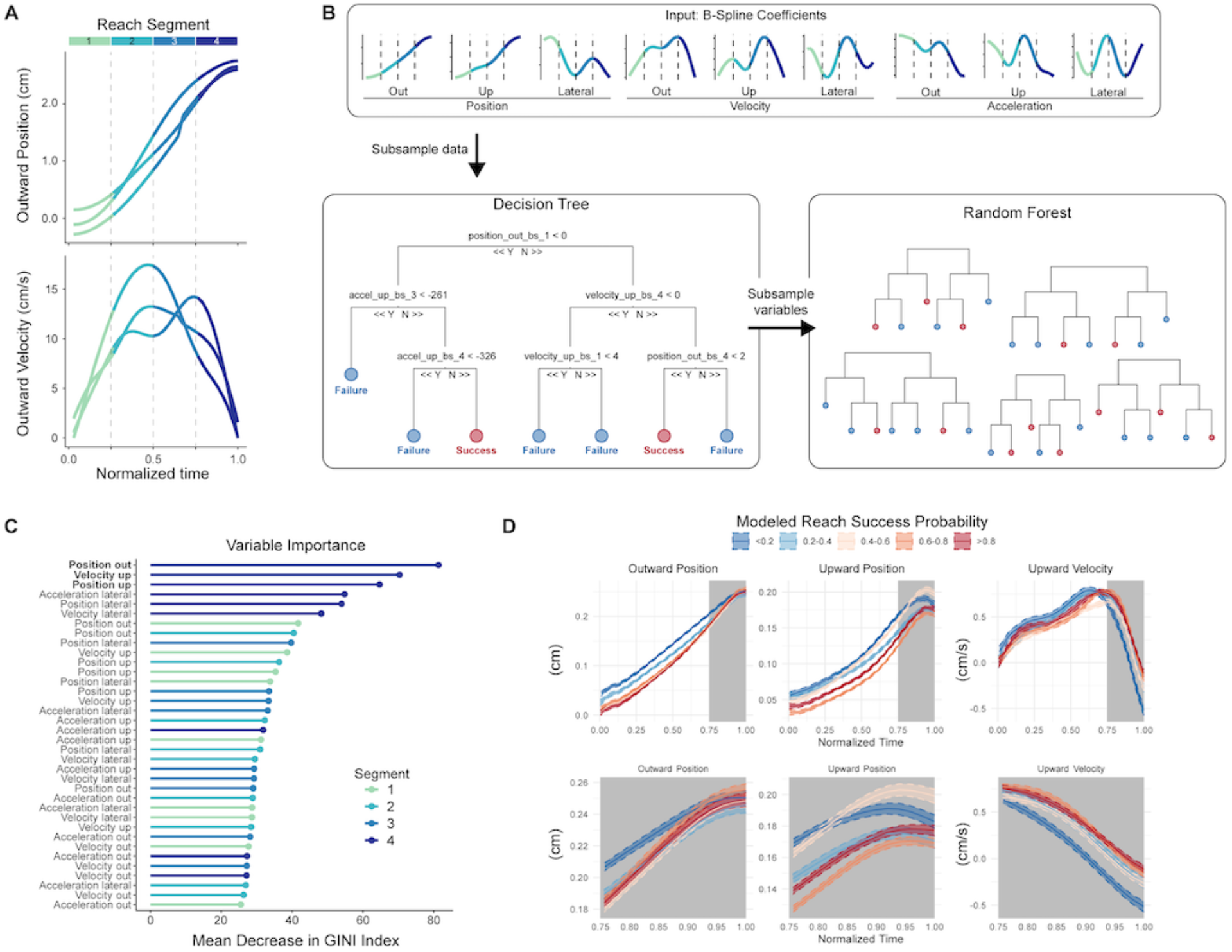
Machine learning model identifies kinematic predictors of reach outcome. A. Outward position and outward velocity profiles for three example reaching movements, displaying initial processing steps for data input into the model. Each reach was decomposed into B-spline coefficients and basis functions; these can be represented as four overlapping time segments (plotted in color). B. Schematic of the random forest model. Variables fed into the model are the four time segments for each of the 9 kinematic variables. Decision trees formed using random subsets of the data split these variables to determine combinations of variable values that predict reach success. Overall results come from aggregating results of a “forest” of trees. C. Ordered plot of variable importance generated by the model for all reaches grouped together. Parameter identity is listed on the y-axis, with time segments plotted in color. High values of mean decrease in GINI index indicate increased importance to model prediction. D. Average kinematic profiles across varying levels of predicted reach success probability (color) by the model. Gray shading indicates the kinematic variable segments that were the top three predictors of success for the model. Top row is full kinematic trajectories, bottom row is last reach segment only.

We implemented the random forest model on data from each mouse individually and on reaches pooled across mice. We first tested how well the model could predict success. Recent work has questioned the relative contribution of ‘gross’ movements (e.g. limb kinematics) versus ‘fine’ movements (e.g. grasping kinematics) in contributing to reach success in trained rodents (Lemke et al., 2019). Even though our data was limited to paw trajectories, kinematic variables accurately predicted reach success for the pooled reaches (Figure 3B, black dots; Training set accuracy: 91%, Test set accuracy: 81%), and for the vast majority (16/17) of individual animals (Figure 3B, gray dots; Training set accuracy: 90.2% +/− 3.7%, range 84-96%; Test set accuracy: 82.1% +/− 8.8%, range 70-94% accuracy). The model’s ability to predict reach outcome suggests that active control of paw kinematics is crucial to motor performance. We next analyzed the importance of each kinematic variable segment to outcome prediction. Since endpoints are significantly more variable on failed reaches, we hypothesized that kinematic variables near reach endpoint would be most informative to outcome.

As predicted, the top 6 most important variables directing prediction of success or failure came from the final (fourth) segment of the reach (Figure 3C). In addition, segments of the position and velocity trajectories tended to be more important than those of acceleration (Figure 3C). If reach trajectories were predetermined, early kinematics would be equally as predictive of reach outcome. Instead, we find evidence for late-phase corrections, either through feedback or other mechanisms, that likely collapse early variability in service of endpoint precision.

To gain further understanding of the kinematic correlates of successful and unsuccessful reaches, we took advantage of the ability of our model to output probability of success on a per reach basis. We plotted full kinematic trajectories of the top 3 most important variables for reaches grouped across quantiles of probability of success (Figure 3D), asking whether reaches with high probability of success or failure displayed different kinematic trajectories. Mean trajectories associated with low probability of success tended to be distal (that is, closer to target) earlier in the reach than trajectories associated with low probability of success (Figure 3D, left and middle panels). As the reaches progressed, positional trajectories converged, suggesting online control mechanisms that adjust for initial kinematic variability. In the upward dimension, reaches predicted to be unsuccessful peaked earlier, implying that coordinated precision in the outward and upward dimensions is associated with success. Interestingly, even though differences in mean trajectories appeared largest for earlier segments of the reach, the model relied more heavily on late-phase kinematics, suggesting that minute differences in the approach to reach endpoint contribute strongly to outcome. Overall, our model and statistical analyses imply that reach kinematics are not predetermined early in the movement. Instead, late-phase alterations in reach kinematics are likely to be central to success outcome due to their ability to correct for initial kinematic variability (Chua & Elliott, 1993; Meyer et al., 1988; K. Novak et al., 2002; Reza Shadmehr et al., 2010).

### Intrinsic kinematic relationships within mouse reaching movements

The neural circuits that produce reaching movements exert exquisite control over the spatiotemporal trajectory of the limb, accurately guiding the paw (or hand) to a specific location in space. Since reach control circuits dynamically generate these kinematic patterns, intrinsic relationships between kinematic variables (i.e. within-reach kinematic correlations) provide a window into potential mechanisms of control (Elliott et al., 2001; Harris & Wolpert, 1998b; J Messier & Kalaska, 1999). As an example, we demonstrated above that both speed and distance are important kinematic features for reach control. In the following sections, we extend our analysis to the relationships between speed, distance, and other variables that define reach kinematics and constrain likely mechanisms of neural control.

Peak reach speed is related to endpoint accuracy (Figure 2), which might be due to the necessity of the paw to slow down near the target to faithfully execute a grasp. We therefore sought to examine how paw speed evolves over time during reach, paying special attention to the decelerative phase from peak speed to endpoint, the segment highlighted as important to reach outcome by our machine learning model. In primate reaches, end-effector speed evolves over time in a characteristic bell shape, rising from rest to maximum speed near the midway timepoint of the reach (Jeannerod, 1984; Morasso, 1981; Soechting & Lacquaniti, 1981). These dynamics can be modeled by minimizing jerk (the third derivative of position), a model that assumes no mid-flight feedback corrections, and thus produces straight spatial trajectories (Atkeson & Hollerbach, 1985; Harris & Wolpert, 1998a; Jeannerod, 1984). Comparing the kinematics of empirical reaches to their minimum jerk approximations can provide insight into the magnitude, timing, and directions of online control mechanisms.

We therefore assessed the shape of mouse reach speed profiles, using similar techniques to primate studies. Time-averaged speed profiles were bell-shaped, in accordance with primate reaches (Figure 4A,B). Since different animals reached with different peak speeds, we normalized the speed profiles to peak, as has been shown in primates to reduce variability (Figure 3C; (Atkeson & Hollerbach, 1985; Jeannerod, 1984)). We applied the same procedure to reaches modeled by minimum jerk, allowing for direct comparisons between empirical and model data (Figure 4C). Compared to the minimum jerk model, mouse reaches moved significantly faster early and late in the movement (Figure 4C). To analyze these differences more concretely, we calculated the area under the curve (AUC) for each half of the speed profile on either side of peak. These areas represent the pathlength, which for the minimum jerk model is equivalent to distance traveled, but is higher for mouse reaches due to their spatial curvature. We calculated an asymmetry metric, defined as the fraction of total AUC that occurred in the first half of the reach, with values less than 0.5 indicating skew towards the decelerative phase (Figure 4D). Mouse speed profiles were asymmetric towards the second half of the movement, again highlighting late-phase alterations in reach kinematics (Figure 4E). In primates, the speed profiles of short distance reaching movements are more symmetric than those of long reaches, which also tend to be skewed toward the second half of the reach (K. Novak et al., 2002). We tested whether this relationship held true for mice by sorting reaches based on their distance travelled and measuring the asymmetry of the speed profile. Reaches in the bottom quartile of distance were comparatively most symmetric (perfect symmetry = 0.50), with symmetry decreasing as a function of distance (Figure 4E; Top: 0.48 +/− 0.02; Bottom: 0.46 +/− 0.02; paired t-test p = 0.0002). The late-phase deviations were especially prominent, with larger differences from minimum jerk occurring in the second half compared to the first half of the reach (Figure 4F; Difference in speed, empirical – model; first half, 2.4 +/− 0.6 cm/s; second half, 3.8 +/− 0.8 cm/s; Wilcoxon signed-rank, p < 0.0001). Overall, these data suggest that mouse reach speed profiles share strong homology with primate reaches, and point to the decelerative phase of the reach as a likely locus for feedback-based or mid-flight adjustment of kinematics.

**Figure 4.**
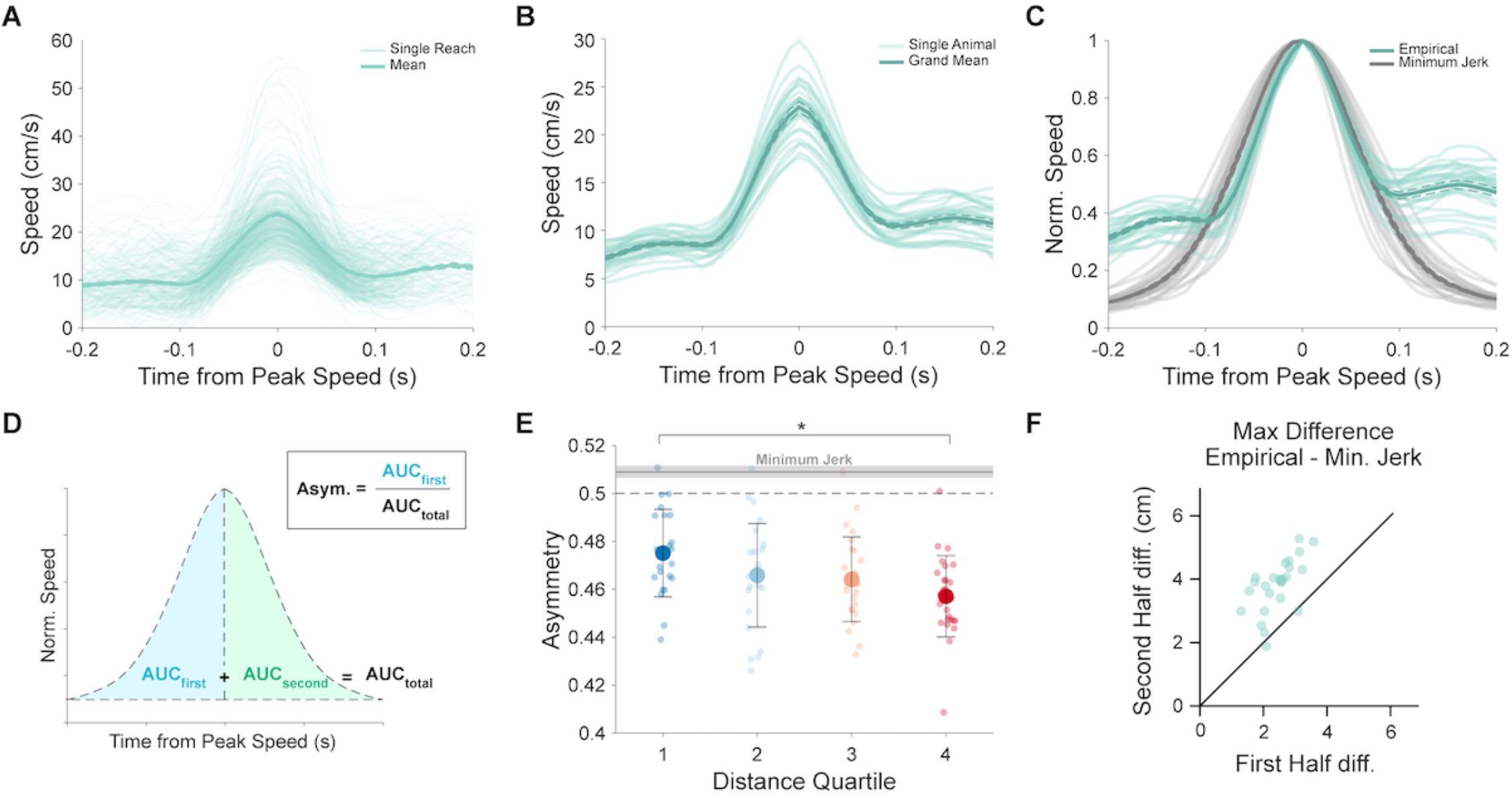
The bell-shaped speed profile of mouse reaches and its relationship to minimum jerk. A. Individual speed versus time profiles (thin lines) and mean speed profile across reaches (thick line) aligned to time of peak speed for an example animal. Average is plotted as mean +/− standard error (dashed lines). B. Individual animal speed profiles (light) and grand mean speed profile (dark), aligned to time of peak speed. C. Empirical individual animal speed profiles (light green) and grand mean speed profile (dark green) aligned to time of peak speed, and normalized to peak speed on a per animal basis. Modeled speed profiles according to minimum jerk per animal (light gray) and mean across animals (dark gray). D. Schematic description of Asymmetry calculation, which involves dividing the area under the curve (AUC) of the first half of the speed profile by the total AUC. E. Individual animal (small points) and grand mean (large points) speed profile asymmetry for reaches grouped into quartiles according to reach distance. Asymmetry is calculated as depicted in D. Mean (gray line) +/− standard deviation (gray shading) asymmetry of minimum jerk modeled reaches. Note that minimum jerk modeled reaches are slightly asymmetric towards the first half of the reach due to subsampling matched to empirical data (see Methods). F. Maximum difference between empirical and minimum jerk speed profiles for the first half (x-axis) and second half (y-axis) of the profile for each animal.

Reach speed and distance, when taken together, implicate a third kinematic variable: duration. For example, given a reach of a specific speed, acquiring targets at different distances will take different amounts of time. We therefore investigated the interrelationships of these thee kinematic variables on a per reach basis. In primates, reaches to more distant targets have higher peak speeds (Jeannerod, 1984). Mouse reaches showed a similar significant positive correlation between distance and peak speed (Figure 5A,B; mean r = 0.24; 24/26 animals p < 0.05). A shuffle control showed no significant relationship (Figure 5B, right; mean r = 0.00; see Methods). In addition, reaches in the top distance quartile had higher peak speeds on average than those in the bottom distance quartile (Figure 5C; Top: 25.7 +/− 3.2 cm/s; Bottom 20.8 +/− 3.0 cm/s; paired t-test, p < 0.0001). If reaches that travel farther have higher peak speeds, there may be a relatively weaker relationship between reach duration and distance travelled, which has been observed in human reaches (Jeannerod, 1984). However, mouse reach distance was more strongly correlated with reach duration than with peak speed (Figure 5D,E; mean r = 0.58; 26/26 animals p < 0.05). Moreover, the mean durations of the top and bottom quartiles of reach distance were significantly different (Figure 5F; Top: 410 +/− 67 ms; Bottom 215 +/− 44 ms; paired t-test, p < 0.0001; shuffle control mean r = 0.00). Overall, both mouse reaches and primate reaches show positive distance-speed and distance-duration relationships, although the relative magnitude of these relationships is reversed.

**Figure 5.**
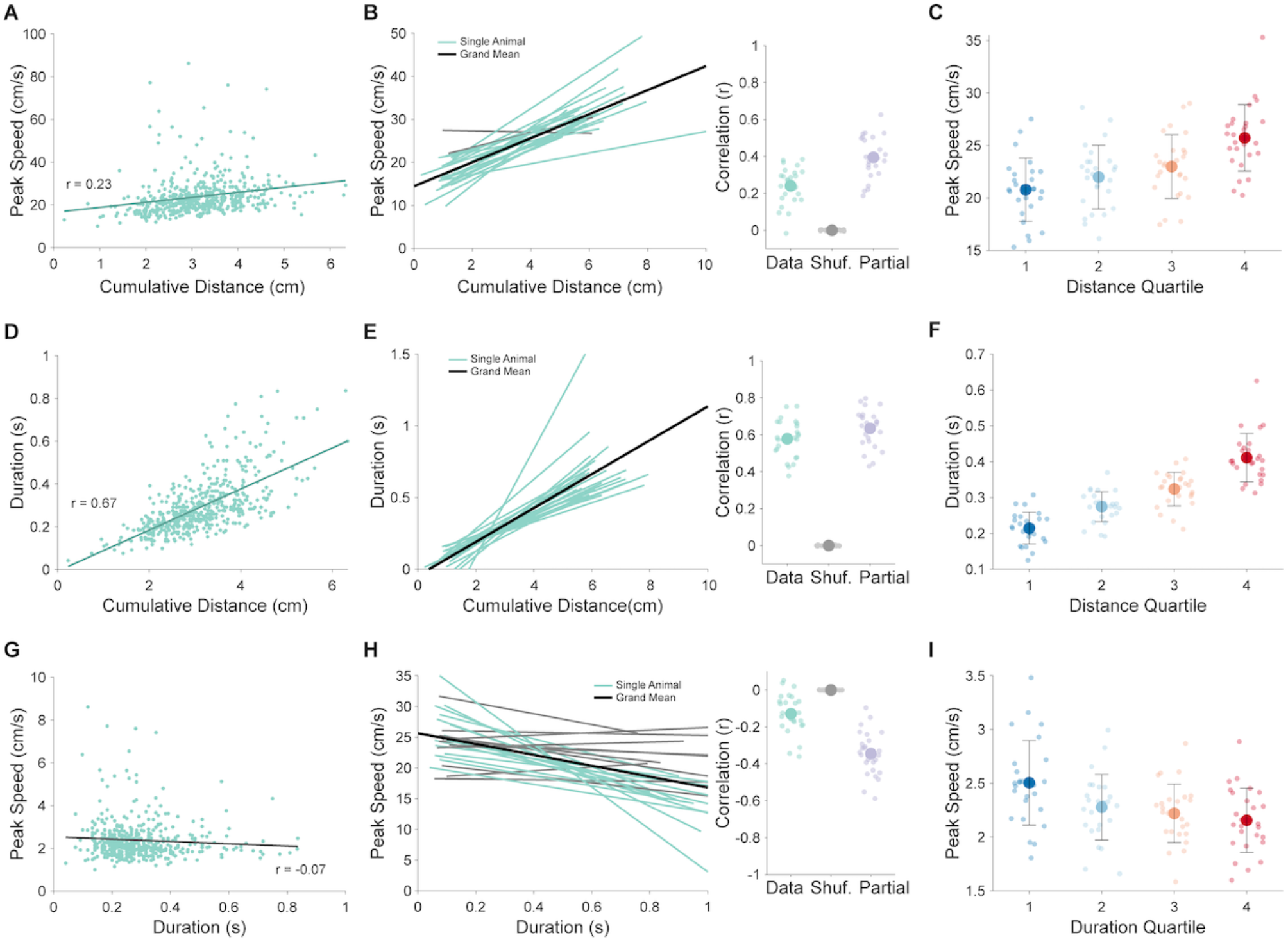
Intrinsic relationships of peak speed, distance, and duration for mouse reach. A. Correlation between cumulative reach distance and peak speed for an example animal. Values from individual reaches (points) are plotted with the line of best fit resulting from simple linear regression. B. Left: Individual animal (thin lines) and grand mean (black) lines of best fit from simple linear regressions relating reach distance to peak speed. Animals displaying significant correlations (p < 0.05) are plotted in blue, with others plotted in gray. Right: per animal correlation values (r) between distance and peak speed, shuffled controls, and partial correlation controlling for duration. C. Individual animal (small points) and grand mean (large points) peak speed values for reaches grouped into quartiles according to reach distance. D. Correlation between cumulative reach distance and duration for an example animal. Values from individual reaches (points) are plotted with the line of best fit resulting from simple linear regression. E. Left: Individual animal (thin lines) and grand mean (black) lines of best fit from simple linear regressions relating reach distance to duration. Animals displaying significant correlations (p < 0.05) are plotted in blue, with others plotted in gray. Right: per animal correlation values (r) between distance and duration, shuffled controls, and partial correlation controlling for peak speed. F. Individual animal (small points) and grand mean (large points) duration values for reaches grouped into quartiles according to reach distance. G. Correlation between reach duration and peak speed for an example animal. Values from individual reaches (points) are plotted with the line of best fit resulting from simple linear regression. H. Left: Individual animal (thin lines) and grand mean (black) lines of best fit from simple linear regressions relating reach duration to peak speed. Animals displaying significant correlations (p < 0.05) are plotted in blue, with others plotted in gray. Right: per animal correlation values (r) between peak speed and duration, shuffled controls, and partial correlation controlling for distance. I. Individual animal (small points) and grand mean (large points) peak speed values for reaches grouped into quartiles according to reach duration.

Intuitively, the correlations described above suggest that since the farthest reaches tend to have higher peak speeds and longer durations, reach duration may be correlated with peak speed. However, from a physical standpoint, reaches that travel faster to the same target must take less time. Indeed, primate reaches appear to lack a discernable relationship between reach duration and peak speed (Jeannerod, 1984). We tested this for mice, and, in contrast to the primate literature, found a weak negative relationship (Figure 5G,H; mean r = −0.13; 15/26 animals p < 0.05). The reaches in the top quartile of duration had relatively slower peak speeds than reaches in the bottom quartile of duration (Figure 5I; Top: 21.6 +/− 3.0 cm/s; Bottom 25.1 +/− 3.9 cm/s; paired t-test, p < 0.0001). These results predict an opposition of peak speed on the distance-duration relationship. We tested this directly by calculating the partial correlations between these three variables. As predicted, accounting for peak speed strengthened the distance-duration relationship (mean partial r = 0.63; paired t-test, Fisher r-to-z, p < 0.0001). Likewise, the partial correlation between distance and peak speed (controlling for duration) also increased (mean partial r = 0.39; paired t-test, Fisher r-to-z, p < 0.0001). Finally, although peak speed and duration were weakly correlated, controlling for distance revealed a significant negative relationship across all animals (mean partial r = −0.35; 26/26 animals p < 0.05; paired t-test, Fisher r-to-z, p < 0.0001). Overall, these data imply strong dependence of mouse reach kinematics on the duration of the reach, emphasizing the importance of neural signals that contribute to reach cessation (J Messier & Kalaska, 1999).

Previous studies, as well as the data presented here, highlight the decelerative phase of reaching movements as particularly important for adjusting to initial variability in reach kinematics, a process that is under direct neural control (Becker & Person, 2019; K. Novak et al., 2002). We therefore investigated how peak velocity was related to subsequent deceleration across the population of reaches in our dataset. By convention, we define the outward direction as positive, meaning deceleration towards the target is represented by a negative value. We predicted a tight relationship between peak velocity and deceleration, as faster reaches necessarily have a larger net change in velocity from peak velocity until endpoint (velocity = 0). As expected, we found that peak outward velocity was inversely correlated with mean deceleration (Figure 6A,B; mean r = −0.46; 25/26 animals p < 0.05), with the fastest quartile of reaches having significantly larger decelerations (Figure 6C; Top: −188.9 +/− 62.5 cm/s2; Bottom: −102.7 +/− 28.4 cm/s2; Wilcoxon signed-rank, p < 0.0001). Nevertheless, the moderate variance explained (R^2^ = .21) implies that other factors must also influence mean deceleration following a given peak velocity. We reasoned that since the goal of a reach is to terminate at a specific target, the position of the limb at peak velocity could influence the mean deceleration necessary to achieve reach accuracy. We therefore sorted reaches by their position of peak outward velocity (PPV), calculating the correlation between peak outward velocity and mean deceleration for each PPV quartile. We found that separating reaches by PPV uniformly resulted in higher correlations between peak velocity and mean deceleration (Figure 6D; mean r; Top: −0.69; Middle-high: −0.73; Middle-low: −0.70; Bottom: −0.60; Fisher r-to-z, paired t-test, p < 0.0001). This result implies that the position of peak velocity accounts for some variability in the peak velocity–mean deceleration relationship. To quantify this effect, we took the partial correlation of peak velocity and mean deceleration, accounting for PPV, and found a significant increase (mean r = −0.67; paired t-test, Fisher r-to-z, p < 0.0001; mean R^2^ = 0.45). Interestingly, we also noticed that the slope of the relationship between peak velocity and mean deceleration changed as a function of PPV. Reaches in the bottom PPV quartile, in which peak velocity occurred further away from the target, had shallower slopes than reaches in the top position quartile (Figure 6E,F; Bottom: −9.5 (cm/s2) / (cm/s); Top: −14.8 (cm/s2) / (cm/s); Wilcoxon singed-rank, p = 0.001). These data imply that decelerative control is informed by both prior velocity as well as the position of the limb. Therefore, any late, decelerative-phase adjustment mechanisms, which are critical for endpoint accuracy, must account for the dynamic interaction between speed and position.

**Figure 6:**
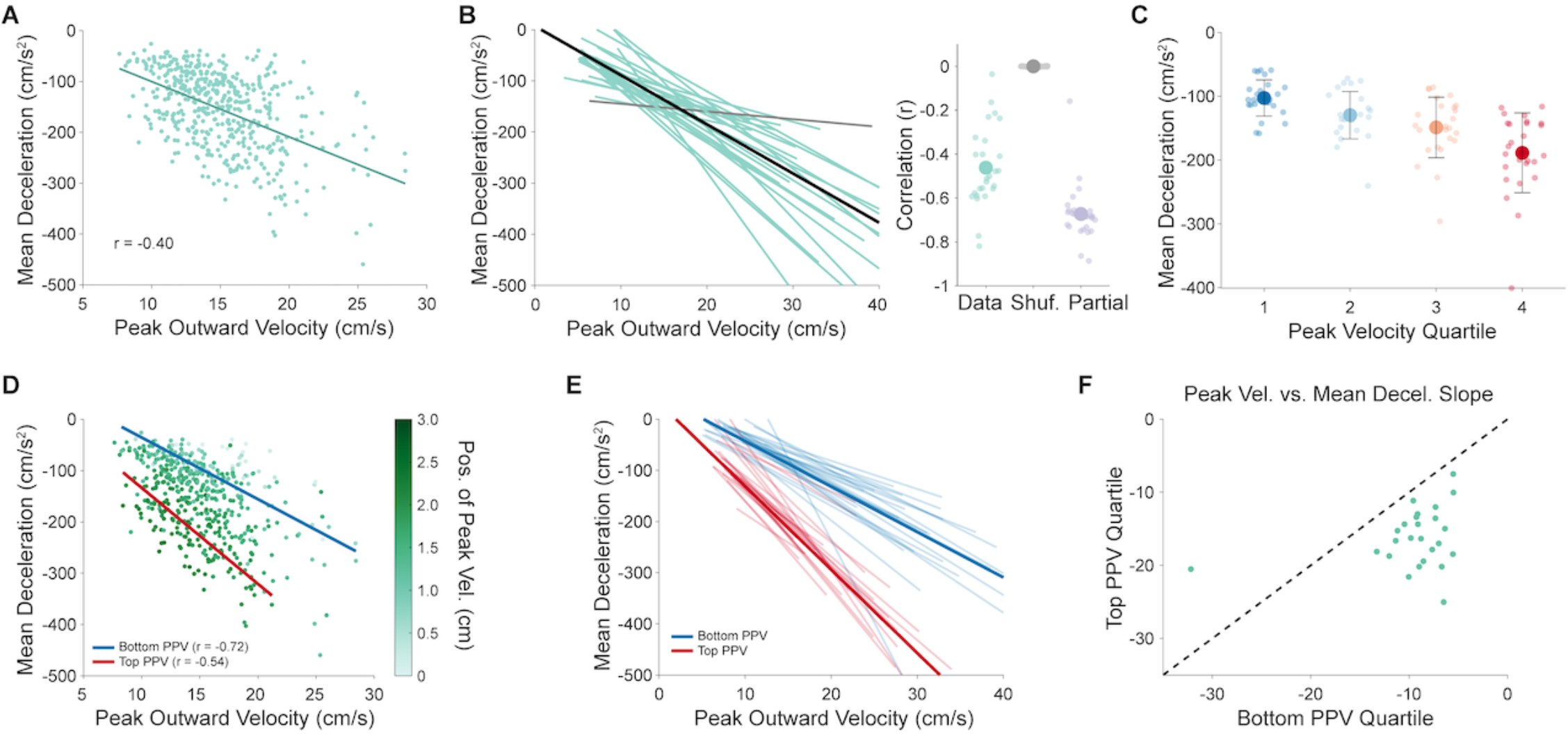
Intrinsic relationships of peak velocity, mean deceleration, and position of peak velocity for mouse reach. A. Correlation between peak outward velocity and mean outward deceleration for an example animal. Values from individual reaches (points) are plotted with the line of best fit resulting from simple linear regression. B. Left: Individual animal (thin lines) and grand mean (black) lines of best fit from simple linear regressions relating peak outward velocity to mean outward deceleration. Animals displaying significant correlations (p < 0.05) are plotted in blue, with others plotted in gray. Right: per animal correlation values (r) between peak velocity and mean deceleration, shuffled controls, and partial correlation controlling for position of peak velocity. C. Individual animal (small points) and grand mean (large points) mean outward deceleration values for reaches grouped into quartiles according to peak outward velocity. D. Same data as in (A) but colored by the position of peak outward velocity (PPV). Lines of best fit from simple linear regressions on data from the top (red) and bottom (blue) quartiles of position of peak outward velocity. The top quartile represents reaches with positions of peak outward velocity that are closer to the target. E. Individual animal (thin lines) and grand mean (thick lines) lines of best fit from simple linear regressions relating peak outward velocity to mean outward deceleration for reaches grouped into the top (red) or bottom (blue) quartiles of position of peak outward velocity (PPV). F. Paired plot comparing individual animal slopes of the lines of best fit from simple linear regressions relating peak outward velocity to mean outward deceleration for reaches in the bottom (x-axis) or top (y-axis) quartile of position of peak outward velocity (PPV). Unity line plotted for reference.

### Trial-over-trial effects on reach accuracy and success

The above analyses consider each reach as a separate event, ignoring their relationship to one another over the course of a behavioral session. Many models of motor learning posit that an error on a given trial can influence the motor plan on the next trial through either explicit (e.g. strategic aiming) or implicit means (e.g. motor adaptation) (Smith & Shadmehr, 2005; Taylor et al., 2014). We tested whether mouse reaches exhibit trial-over-trial error correction by analyzing changes in reach endpoint locations from one reach to the next. Our analysis is rooted in the observation that reaches, as defined by a segmentation algorithm (see Methods), could contain multiple endpoints if the first attempt failed, as the animal continued ‘re-reaching’ to obtain the pellet (Figure 7A). In these trials, we extracted the endpoints of first and second attempts, and analyzed both endpoint location and endpoint variability as two separate tests of error correction. If mean endpoint location is significantly different between attempts, it implies a deficit in accuracy overall, while a change in endpoint variability implies a deficit in precision. Testing endpoint locations from individual animals, we found that first attempts were significantly hypometric in the outward and upward directions compared to second attempts (Figure 7B,C; Outward: 23/26 animals; Upward: 22/26 animals; Wilcoxon signed-rank, p < 0.05). Across the population of animals, first attempts ended short when compared to second attempts (Figure 7D,E; Outward location from origin; First attempts: 2.45 +/− 0.11 cm; Second attempts: 2.56 +/− 0.11 cm; Wilcoxon sign-rank p < 0.0001). In order to quantify whether second attempts are eumetric compared to first attempts (i.e. more accurate), we calculated the mean endpoint of final attempts on all successful trials on a per animal basis. We reasoned that the last endpoints of a successful reach represent the movement segment in which the animal actually obtained the pellet, and thus is a good estimate of accurate hand endpoint positions. We predicted that second attempts might be closer to this ‘target’ location, and thus eumetric, compared to first attempts. As predicted, second attempt endpoints were significantly closer to target than first attempts, in both outward and upward dimensions (Figure 7F; Distance to target [endpoint – Target]; First attempts: −0.09 cm out, −0.16 cm up; Second attempts: 0.02 cm out, −0.05 cm up; Wilcoxon signed-rank p < 0.0001). Even though first reach attempts were hypometric and further from the target, first attempt endpoints were typically not more variable than second attempts within animals or across the population (Figure 7D,E, right panels; 5/26 animals, Levene’s Test, p < 0.05; mean standard deviation, Wilcoxon signed-rank p = 0.30). Overall, these results imply that successive reach attempts are more accurate, as animals correct for their initial hypometria, but that mechanisms controlling reach precision are intact in both initial and subsequent attempts.

**Figure 7:**
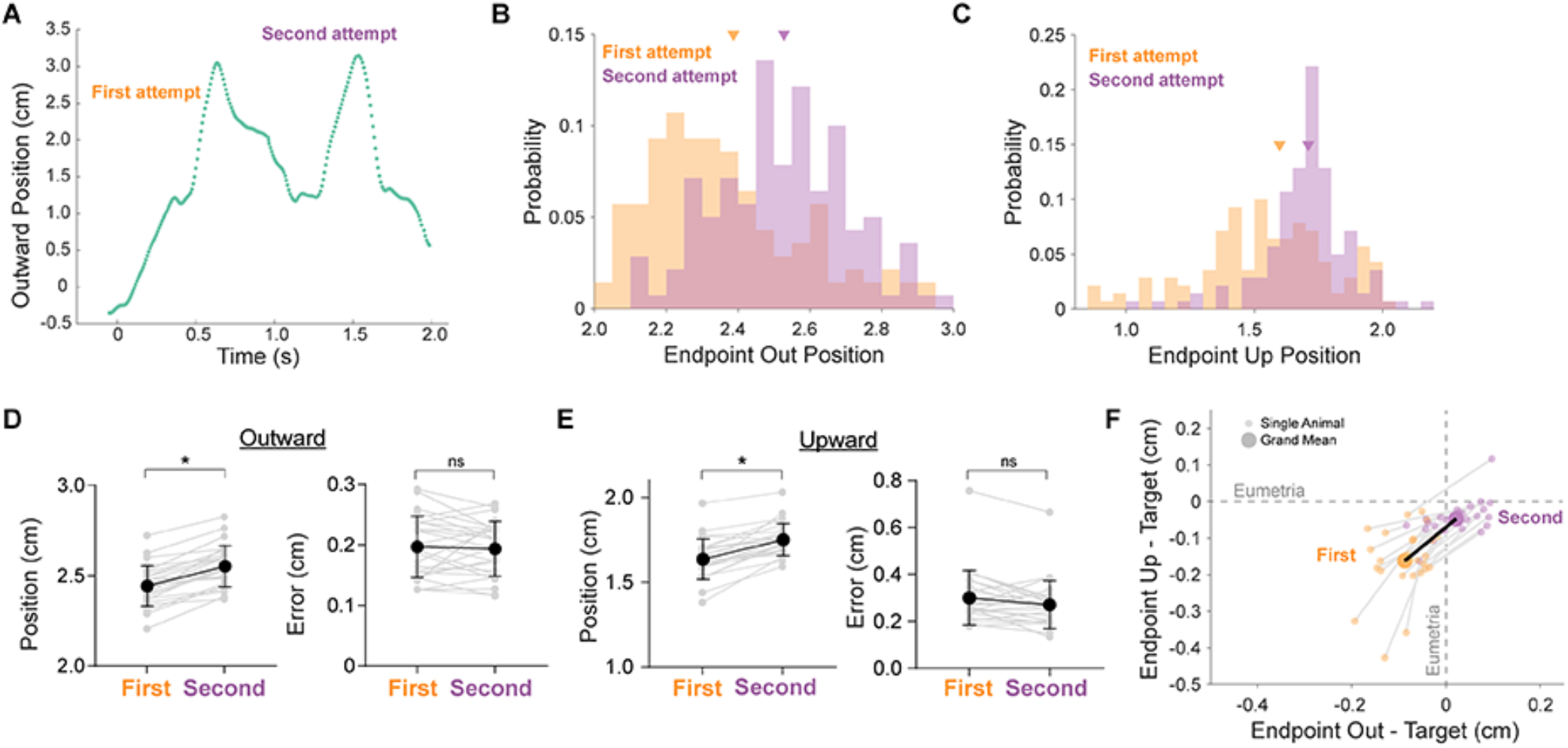
Endpoint statistics of consecutive reach attempts. A. Example reach trajectory in the outward direction versus time, demonstrating a single reach (as identified by our segmentation algorithm, see Methods) that contains multiple attempts. B. Distributions of endpoints in the outward dimension on first (orange) and second (purple) consecutive attempts from an example animal. Group means are indicated by arrowheads. C. Distributions of endpoints in the upward dimension on first (orange) and second (purple) consecutive attempts from an example animal. Group means are indicated by arrowheads. D. Individual animal (gray) and grand mean (black) average endpoint position (left) and standard deviation (right) in the outward dimension for first and second attempts. E. Individual animal (gray) and grand mean (black) average endpoint position (left) and standard deviation (right) in the upward dimension for first and second attempts. F. Individual animal (small points) and grand mean (large points) average endpoint positions relative to target location in the outward and upward dimensions for first attempts (orange) and second attempts (purple). Target location was calculated on a per animal basis as the mean outward and upward endpoint location of final attempts on successful trials. Values of zero represent eumetric endpoints relative to target.

The above results suggest that reach accuracy improves from one reach to the next, as animals home in on the correct target endpoint. If this process occurs across reaches throughout an entire behavioral session, it has two alterative implications. One possibility is that reach accuracy is maintained once it is achieved, resulting in continually successful outcomes once the target location has been obtained. On the other hand, motor variability may cause a recurrence of inaccurate reaches, which, outside of this specific behavioral task, could enable flexibility and motor exploration (Dhawale et al., 2017; Tumer & Brainard, 2007). We therefore sought to extend our trial-over-trial analysis to include the influence of past reach success or failure on subsequent reaches. A famous formulation of this problem is referred to as ‘hot-handedness’: an increase in the likelihood of success given a string of consecutive successes (Daks et al., 2017; Gilovich et al., 1985; Miller & Sanjurjo, 2015). For example, if a mouse successfully obtains a pellet three reaches in a row, is that mouse more likely to obtain a pellet on the subsequent reach? We asked whether the mice in our task exhibited a hot hand by conducting a permutation test (10,000 repetitions) for streaks (n) of 1,2, or 3 successes across all behavioral sessions (Daks et al., 2017). For each session, we calculated the probability of a success given a string of n previous successes, minus the probability of a success given an equally long string of consecutive failures. We then compared this single-session probability with the distribution of probabilities calculated across 10,000 randomly generated permutations of the original data. If the actual session probability fell within the top 5% of the random distribution, we considered it significant at the p < 0.05 level. Of the 30 sessions analyzed, 3 showed significant hot-handedness for a streak of 1; 1 session showed hot handedness for a streak of 2; and no sessions showed hot-handedness for streaks of 3 success. Cold-handedness was similar, with no sessions showing significance for 1 and 3-streaks and only 2 showing significance for increased chance of success after a 2 failure streak. Thus, while reaching behavior in mice displays streakiness in a small minority of behavioral sessions, hot-handedness is rare overall, similar to skilled human motor behaviors.

### Mouse reach kinematics in the head-fixed context

Motor behavior is strongly influenced by the context in which it is performed. Previous work identified qualitative differences in the behavioral patterns of reach-to-grasp movements in mice that were head-fixed, compared to freely-behaving animals, but whether these differences extend to the level of reach kinematics remains unclear (Whishaw et al., 2017). We therefore sought to test the degree to which the intrinsic kinematic relationships defined above are altered by the experimental constraint of head-fixation, a relatively common experimental method (Figure 8A-C). Head-fixed animals (N = 4; mean n = 240, range 113 – 344, total 960 reaches) were trained to perform an identical reach-to-grasp task, including use of the same behavioral training procedure, food pellet targets, motion capture tracking system, and kinematic analyses. We noticed some basic differences between freely-behaving (FB) and head-fixed (HF) reaches (Figure 8). HF reaches were faster on average, including higher peak speeds, outward velocities, and upward velocities (Figure 8 F-H). In addition, the distance and duration of HF reaches trended lower in magnitude and were clearly less variable than FB reaches (Figure 8 D,E), likely due to the constraint on reach starting position under head-fixation.

**Figure 8.**
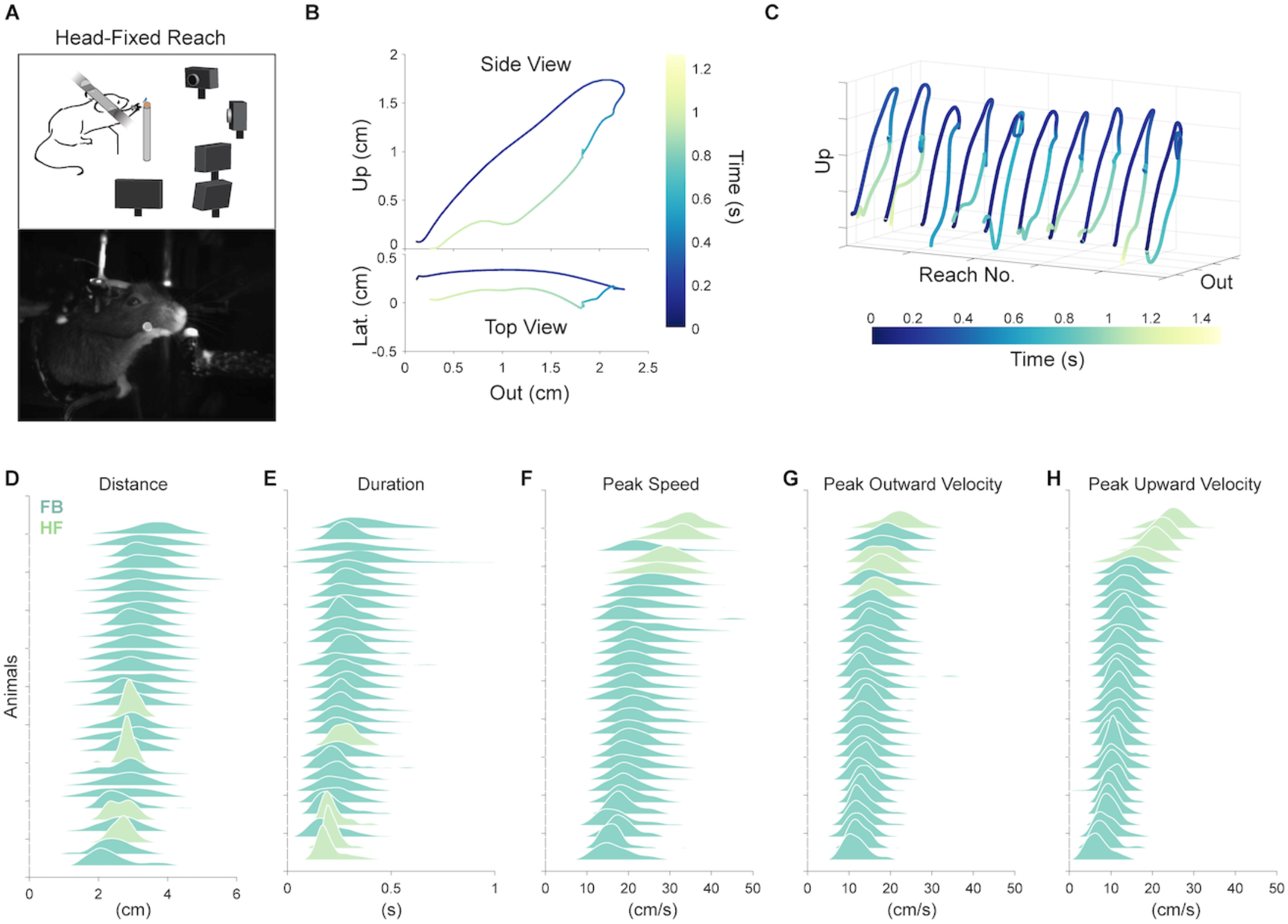
Distribution of basic kinematic parameters of freely-behaving and head-fixed reaches. A. Behavioral setup. Top, schematic of head-fixed reach tracking behavioral arena. The same tracking system is used as in freely-behaving experiments (Figure 1). Bottom, image of a head-fixed mouse performing a reaching movement. B. Example head-fixed mouse reach trajectory. Position is plotted from a side view (top) and top view (bottom) with time indicated in color. C. Several example reach trajectories from a head-fixed mouse. D. Joyplot displaying the distribution of reach distance values for each freely-behaving (FB; teal) and head-fixed (HF; olive) animal. Animals are sorted by mean reach distance. E-H. Same as in A, but for Duration, Peak Speed, Peak Outward Velocity, and Peak Upward Velocity, respectively.

We wondered whether these differences in the distributions of basic reach kinematic parameters would affect the intrinsic kinematic relationships identified above. The relationship between speed, distance, and duration was mostly preserved in HF reaches (Figure 9A,B). We found a moderately stronger correlation between distance and duration, and a weaker correlation between distance and peak speed, in HF reaches compared to FB reaches. A weaker relationship between peak speed and distance implies that mean deceleration may be strongly coupled to peak speed, resulting in reaches of similar distance. We therefore turned to the core relationship between peak velocity, mean deceleration, and position of peak velocity, and similarly found that it was preserved in HF reaches (Figure 9C,D). Peak outward velocity was strongly correlated to mean deceleration, with moderately stronger correlations than in FB animals (mean r; HF: −0.59; FB: −0.46). We hypothesized that the increased strength of the peak velocity-mean deceleration relationship could be due to a decrease in positional variability in the HF condition compared to the FB condition. If this is true, accounting for the position of peak velocity (PPV) should have a relatively modest effect on the relationship between peak velocity and mean deceleration. As predicted, the partial correlation of peak velocity and mean deceleration accounting for PPV was only slightly increased (mean partial r: −0.67). Intriguingly, this mean partial correlation was identical to the partial correlation values of FB reaches (mean partial r = −0.67). In summary, the weaker relationship between peak velocity and mean deceleration in FB reaches can be accounted for by positional variability, resulting in equal amounts of variance in mean deceleration explained (R^2^ = 0.45) in the two conditions when accounting for both position and speed. Overall, HF and FB reaches share several core intrinsic kinematic features, with a reduction in positional variability likely accounting for differing magnitudes of correlations.

**Figure 9.**
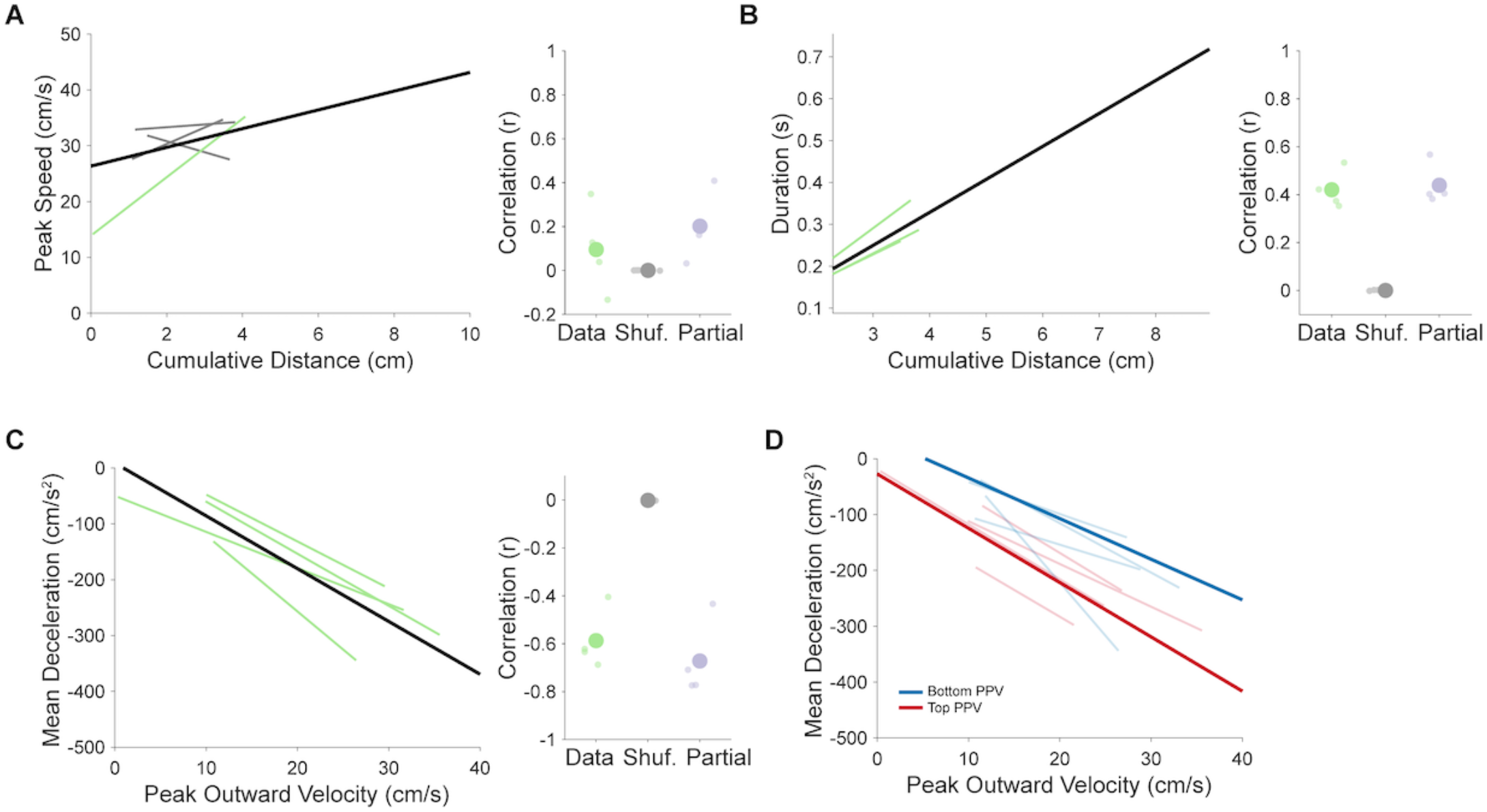
Intrinsic kinematic relationships of head-fixed reaches. A. Left: Individual animal (thin lines) and grand mean (black) lines of best fit from simple linear regressions relating reach distance to peak speed Grand mean regression is plotted over a wide range to facilitate comparison to freely-behaving reaches (Figure 5). Animals displaying significant correlations (p < 0.05) are plotted in green, with others plotted in gray. Right: per animal correlation values (r) between distance and peak speed, shuffled controls, and partial correlation controlling for duration. B. As in A, but comparing reach distance to duration, with partial correlations controlling for peak speed. C. As in A, but comparing peak outward velocity to mean deceleration, with partial correlations controlling for position of peak velocity. D. Individual animal (thin lines) and grand mean (thick lines) lines of best fit from simple linear regressions relating peak outward velocity to mean outward deceleration for reaches grouped into the top (red) or bottom (blue) quartiles of position of peak outward velocity (PPV).

Reach kinematics are intimately tied to outcome. We therefore investigated whether the altered reach kinematics observed for HF reaches led to differences in success or accuracy. Overall, success rates were higher in the HF condition compared to FB (mean success rate: FB, 32.0 +/− 9.9%; HF, 49.2 +/− 5.2%; Wilcoxon rank sum p = 0.0031). High success rates could be due to decreased positional variability of HF animals over the course of a session. However, HF reaches also showed higher peak speeds, which in the FB condition is associated with greater endpoint error. We therefore separated reaches based on peak speed or velocity, as done previously for FB reaches, and measured reach endpoint error in three dimensions, to see if the speed-accuracy tradeoff is preserved under head-fixation. Surprisingly, we found an inverted trend in the relationship between peak speed and endpoint error; the fastest HF reaches showed decreased endpoint error compared to the slowest HF reaches (Figure 10A-C; Speed: Top quartile error 0.53 +/− 0.22 cm, Bottom 0.44 +/− 0.17 cm, Wilcoxon signed-rank p = 0.13; Outward velocity: Top 0.57 +/− 0.26 cm, Bottom 0.43 +/− 0.18 cm, p = 0.13; Upward velocity: Top 0.59 +/− 0.24 cm, Bottom 0.39 +/− 0.17 cm, p = 0.13). This result led us to investigate the speed and velocity profiles of HF reaches themselves. The speed profile of HF reaches was highly asymmetric, compared both to the minimum jerk model as well as the FB reaches (Figure 10D; see Figure 4). Decomposing speed into outward and upward velocity profiles revealed that the upward profile in particular was asymmetric (Figure 10E,F). In HF reaches, upward velocity typically peaked at more proximal positions than outward velocity, while in FB reaches, positions of peak upward and outward velocity largely overlapped or showed the opposite relationship (Figure 10G,H; mean PPV Out – PPV Up: FB, −0.20 +/− 0.22 cm; HF, 0.42 +/− 0.26; Wilcoxon rank sum p = 0.003). The decoupling of upward and outward velocity in HF reaches could reflect an altered motor strategy for obtaining the pellet in the HF condition compared to the FB condition, where outward and upward dimensions are highly coordinated. Overall, analysis of HF and FB reaches reveals that shared core kinematic relationships that can be altered by the physical constraints of highly similar tasks, likely due to shifts in strategy.

**Figure 10.**
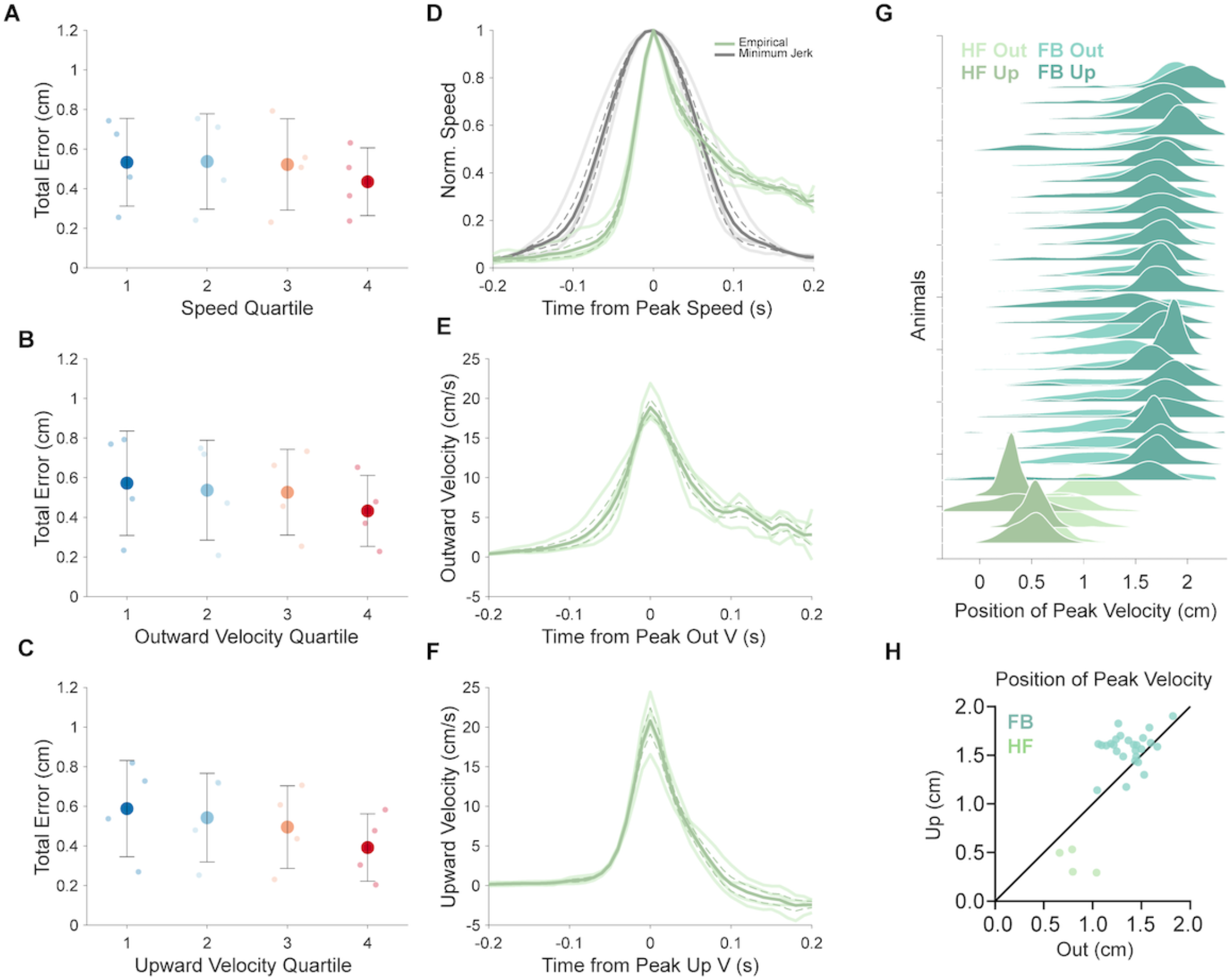
Head-fixed reach speed-accuracy relationship and velocity profiles. A. Individual animal (small points) and grand mean (large points) total error of endpoints for reaches grouped into quartiles according to peak speed. Reaches were grouped on a per-animal basis. Total error is defined as the sum of the standard deviations calculated individually for each positional dimension (outward, upward, lateral). B, C. Individual animal (small points) and grand mean (large points) total error of endpoints for reaches grouped into quartiles according to peak outward velocity (B), peak upward velocity (C). D. Empirical individual animal speed profiles (light green) and grand mean speed profile (dark green) aligned to time of peak speed, and normalized to peak speed on a per animal basis. Modeled speed profiles according to minimum jerk per animal (light gray) and mean across animals (dark gray). E. Outward velocity profiles aligned to time of peak outward velocity. Convention as in D. F. Upward velocity profiles aligned to time of peak upward velocity. Convention as in D. G. Joyplot comparing position of peak velocity in the outward (light) and upward (dark) directions between freely-behaving (teal) and head-fixed (olive) conditions. Sorted by mean position of peak outward velocity. H. Comparison of per-animal mean position of peak velocity in the outward (x-axis) and the upward (y-axis) directions for freely-behaving (teal) and head-fixed (olive) reaches.

## Discussion

In this study, we establish a set of quantitative kinematic observations that constrain the possible neural mechanisms underlying skilled reaching movements. Our analyses focus on endpoint and the factors that directly regulate its location, highlighting the decelerative phase of reach as an especially important time window for active control. Both explicit comparisons to a well-established movement trajectory model (minimum jerk) as well as an unbiased machine learning prediction model directly implicate the late-phase of reach as a site of kinematic adjustment. In turn, we find that mean deceleration is linked not only to peak velocity, but the position of the limb at peak velocity, implying that reach control systems must integrate diverse information streams in their influence on endpoint accuracy. Analyzing successive reach attempts, we find evidence for an endpoint error correction process, which stands in contrast to a lack of a trial-over-trial effect of outcome on subsequent performance. Finally, comparison of freely-behaving and head-fixed reach kinematics provides a clear example of how task constraints can alter motor control processes, but also identifies kinematic relationships that are stable across contexts and across species.

### Comparative kinematics support reach homology across species

Throughout this study, our goal was to bridge the gap between prominent quantitative kinematic frameworks developed in primate models and the relatively nascent neural-mechanistic studies of mouse skilled forelimb control. The use of powerful genetic and molecular technologies in mice to understand neural contributions to behavior is inherently limited by the degree to which the behavior itself is measured, quantified, and understood from a theoretical and algorithmic perspective (Krakauer et al., 2017). Thus, charting similarities and differences in basic kinematic measurements across behavioral contexts and across species can provide stepping stones for future experiments aimed at uncovering neural mechanisms into how the nervous system produces reach behavior. Although some prior studies quantified aspects of mouse reach kinematics, the majority focused mainly on success rates, qualitative descriptions and scoring systems, or categorical demarcation of the reach into phases, with few explicit connections drawn to the kinematic control principles discussed here (Esposito et al., 2014; Farr & Whishaw, 2002; Galiñanes et al., 2018b; Guo et al., 2015; Whishaw, 1996; Whishaw et al., 2017, 2018; Whishaw & Pellis, 1990). This study establishes several basic quantitative frameworks for analyzing mouse reach kinematics, allowing numerical comparisons to be drawn across species and across tasks. Moreover, in analyzing thousands of trials, we identify robust correlational features across mice and across the highly-variable motor behavior of skilled reach.

Consequently, we identified strong homology to primate reach kinematics across a variety of analyses. Shared kinematic features of mouse and primate reaches include the speed-accuracy tradeoff, bell-shaped velocity profile (including late-phase asymmetry), the ‘main sequence’ correlation of speed, distance, and duration, and slightly hypometric endpoints on failures (Chua & Elliott, 1993; Fitts, 1954; Harris & Wolpert, 2006; Jeannerod, 1984; Morasso, 1981). These results lend support to the idea of a shared evolutionary origin for skilled forelimb movements early in the tetrapod lineage (Iwaniuk & Whishaw, 2000; Whishaw et al., 1992). While basic kinematic relationships were conserved, some notable differences were found as well. For example, mouse reaches displayed a relatively stronger relationship between distance and duration, rather than distance and peak speed (Jeannerod, 1984). The relative strength of these intrinsic kinematic relationships could be influenced by the nature of the task, which can dictate strategy. Concomitantly, we found task-dependent alterations in the speed-accuracy tradeoff, with head-fixed reaches displaying greater endpoint error on the slowest instead of the fastest reaches (Dounskaia et al., 2005; Fernandez & Bootsma, 2004; Roberts et al., 2016). Compared to the freely-behaving condition, head-fixed reaches displayed an early upward velocity peak that dominated the speed profile, resulting in high asymmetry. If early peaks in upward velocity correspond to an initial ‘aiming’ phase separate from the ‘transport’ phase, it could represent an altered control strategy interacting with the constraints of head-fixation (Whishaw et al., 1992, 2017). Ultimately, these similarities and differences in basic kinematic features represent compelling avenues for further exploration of species- and task-specific constraints on reach control.

### Kinematic evidence for the importance of late-phase control

Despite the nearly-infinite possible trajectories for a given reach target, canonical principles of kinematic organization exist, with direct implications for underlying mechanisms of neural control (Bernstein, 1967; Lashley, 1933; Latash, 2012; Li, 2006). A common underlying principle tying reach kinematics to neural systems is the interaction of signal-dependent noise with speed (i.e. the speed-accuracy tradeoff), which may contribute to bell-shaped velocity profiles as well as the ‘main sequence’ correlational structure (Harris & Wolpert, 1998b, 2006; Meyer et al., 1988). In general, reaching movements are typically straight and smooth and can be simply modeled by minimizing jerk (Atkeson & Hollerbach, 1985; Harris & Wolpert, 1998b; Jeannerod, 1984). However, since the minimum jerk model does not account for motor feedback or noise, deviations can be interpreted as evidence for feedback control (Fernandez & Bootsma, 2004). Mouse reaches consistently displayed bell-shaped speed profiles that deviated from minimum jerk most prominently during the decelerative phase of the reach, implying late-phase adjustments of reach kinematics, presumably via feedback corrective mechanisms. Using submovement decomposition, previous work showed that late asymmetries in the speed profile likely represent small ‘submovements’ that are added on to the end of the reach, enhancing reach accuracy (K. Novak et al., 2002; K. E. Novak et al., 2003). We observed that reaches of shorter distances were more symmetric, implying a greater utilization of these late-phase adjustments during reaches that travel farther. However, we also demonstrated that more distant reaches generally have higher peak velocities, and that accuracy is decreased in the fastest reaches. This could be viewed as an apparent contradiction: the hallmarks of late-phase adjustments are more prevalent in more distant reaches, but the reaches remain less accurate. On the contrary, we argue that late-phase velocity asymmetries represent an adaptive response to the increased kinematic variability of distant (and hence faster) reaches. In our view, the inherent tradeoff between speed and accuracy could be mitigated by a shared neural mechanism of online kinematic control that adjusts for initial variability in service of endpoint precision (Becker & Person, 2019; J Messier & Kalaska, 1999). Without this neural mechanism intact, we predict a decrease in reach precision, potentially resembling the oscillatory dysmetria of cerebellar patients (Diener & Dichgans, 1992; Hore et al., 1991).

In addition to these classical methods, we created a random forest model to identify and rank kinematic predictors of reach outcome. In essence, our model allowed us to ‘screen’ kinematic parameters, including their progression throughout the reach, for their contribution to success, thus identifying task-relevant variables. Again, our model highlighted the late-phase of reach, particularly position and velocity, as central to performance. Interestingly, even though the random forest model inputs were limited to end-effector tracking data, not accounting for digit kinematics, the model was able to predict success with a high degree of accuracy. Taken together, our use of explicit and machine learning models provides strong evidence for active control processes late in the reach that alter kinematics to produce successful pellet retrieval.

We therefore turned to the decelerative phase of reach specifically, identifying a key relationship between peak velocity, mean deceleration, and position of the limb at peak velocity. As implied by deviations from minimum jerk models and asymmetric velocity profiles described above, we found a moderate positive relationship between peak speed and mean deceleration in freely-behaving reaches, which was significantly improved when accounting for the position of peak velocity. If mean deceleration is influenced by late-phase corrective submovements, it follows that submovement amplitude could depend on the position of the limb at peak velocity. In this way, the relationship between position and velocity could be used as a substrate for predicting kinematic corrections needed to slow the limb accurately to target. Prediction is essential here, as the delays associated with sensory processing limit the utility of purely feedback-based mechanisms. Interestingly, the peak speed-mean deceleration relationship showed less reliance on position in head-fixed reaches, implying a degree of context-specific flexibility in the late-phase adjustments that sculpt the decelerative profile. The degree to which intrinsic kinematic relationships serve as substrates for motor learning, or reflect the outcome of previously learned motor adjustments, remains an open question.

### Regulation of motor variability

Though our analysis was limited to reaches performed in the learned state, we were interested in identifying control processes that operate from reach-to-reach throughout a behavioral session. First, we found that reach endpoint appears to be actively regulated in trials in which animals make several attempts during a single reach. A previous study reported multiple-attempt reaches in head-fixed mice, describing a touch-release-grab sequence in which the initial attempt involves “semi-flexed and semi-closed digits” (Whishaw et al., 2017). We extended this observation by analyzing endpoints on subsequent reach attempts, finding that the mean distance from the target decreased from first to second attempts with no significant difference in endpoint variability. These results are consistent with the animal touching the front of the food pedestal on the first attempt, then reaching to grasp the pellet on the second attempt. The pattern of multiple attempts that ‘home in’ on the target is seen in reaching movements of human subjects in the dark (Chua & Elliott, 1993; Khan & Franks, 2003; Spijkers & Lochner, 1994) and tongue movements of mice licking for water, suggesting a broad neural mechanism in control of various end-effectors (Bollu, Whitehead, Kardon, et al., 2019).

The observation that endpoint variability is equivalent on first and second attempts implicates a neural mechanism that adjusts endpoint positioning with no effect on overall precision. Determining if this pattern is based on sensory feedback mechanisms within a specific trial, or is an overall motor control strategy unrelated to feedback, could inform hypotheses about the neural circuits involved in its control. Mice may have difficulty sensing pellet depth; olfactory cues may be somewhat noisy, visual cues appear unnecessary, and somatosensation is precluded as whiskers are blocked from the target (Galiñanes et al., 2018a; Whishaw & Tomie, 1989). Importantly, we observed multiple attempt reaches in both freely-behaving and head-fixed conditions, pointing to a broader reliance on the multiple-attempt strategy than previously suggested. When we analyzed the effect of endpoint position on outcome, we found the opposite result: unsuccessful reaches terminated slightly hypometric to the target in some animals, but accuracy effects were overshadowed by large differences in endpoint variability. These results suggest that neural mechanisms reducing variability increase performance, as observed in both humans and animals, and are applied evenly across attempts (Dhawale et al., 2019; Shmuelof et al., 2012; Wu et al., 2014).

If variability is reduced indefinitely as a skill is refined, learning new contingencies may suffer, as motor exploration can be critical to adaptation (Churchland et al., 2006; Dhawale et al., 2017, 2019; Tumer & Brainard, 2007; Wu et al., 2014). Consistent improvement over reaches might predict that once a mouse finds a successful strategy it will cease making errors, which would resemble a ‘hot hand’ as reaches continually succeed. We therefore tested the hot-hand hypothesis, which originates from the widespread notion that basketball players that have scored on several consecutive attempts are more likely to score on their next attempt. This notion has received significant attention over the past several decades, mostly as an example of fallacious human reasoning (Gilovich et al., 1985). However, recent disagreements about the statistical assumptions in calculating hot-handedness have resulted in a revision of this view, with researchers finding a small but significant effect of hot-handedness on success outcome (Miller & Sanjurjo, 2015). In an investigation of the 2016-2017 Golden State Warriors, an exceptionally skilled group of basketball shooters, researchers found evidence for a hot-hand in approximately 5% of games, strikingly similar to the frequency of hot-handedness calculated here for mouse reaching (Daks et al., 2017). Together, these observations position mouse reaching behavior as a motor skill, commonly failing even in the learned state similar to basketball shots. More importantly, our results pit task relevant variability against mechanisms that enhance precision, suggesting multiple neural substrates working cooperatively and/or competitively to promote successful motor behavior while remaining adaptable.

## Materials and Methods

### Experimental model

Behavioral data were collected from 30 adult (P70-P365) mice of either sex (27 females, 3 males), including the following genotypes: (1) ‘wild-type’ C57/Bl6 (N = 24); (2) Neurotensin receptor1-Cre (N = 3; Mutant Mouse Regional Resource Center; Tg(Ntsr1-cre)GN220Gsat/ Mmucd) (3) Neurotensin receptor1-Cre x FLEX-ChR2 (N = 3; Flex-ChR2: RCL-ChR2(H134R)/EYFP; Jackson Labs Stock No: 024109). There was no a priori expectation of genotypic or sex differences in reach motor control, thus the study was not powered to examine such potential differences. Animals were housed on a 12:12 light-dark cycle with ad libitum access to food and water except during behavioral training and experimentation (described below), and maintained bodyweights above 80% for the entirety of the study. Animals were group housed with like genotypes until surgery, at which point they were singly housed with a running wheel. All procedures were in accordance with NIH guidelines for the care and use of laboratory animals and were approved by the University of Colorado Anschutz Institutional Animal Care and Use Committee and Institutional Biosafety Committee.

### Kinematic tracking system

Paw position was monitored by an infrared-based real-time motion-capture system described previously (Becker & Person, 2019). Briefly, five 120 frames-per-second cameras (Optitrack Slim3U; Corvallis, OR), each with a 10 mm focal length lens (Edmund Optics, Barrington, NJ) and infrared LED light ring array, were fixed in place with custom built mounts and focused onto a capture volume of approximately 4 cm^3^, large enough to capture the entirety of a mouse reaching movement. Four cameras were used for online kinematic tracking, and one was used for reference video. 1.5 mm diameter retroreflective markers (B&L Engineering, Santa Ana, CA) were used for both camera calibration and tracking of mouse reach kinematics. Camera calibration was conducted in Optitrack Motive software with a custom-built calibration wand (4 and 8 mm marker spacing) and ground plane (10 and 15 mm marker spacing). The spatial origin of the freely-behaving reach capture volume was located approximately 1.6 cm inside the behavioral arena, measured from the front plexiglass wall, and approximately 2.0 mm left-of-center from the reach opening (in line with the pellet target location). The front wall of the behavioral arena defined the ‘upward’ and ‘lateral’ dimensions, with the ‘outward’ dimension defined perpendicularly from that plane toward the pellet. For the head-fixed condition, the origin was set to the center of the bar that mice rest their hands on, with the lateral dimension running parallel to the bar. Any reported positional measurements are in reference to the origin location. Calibration procedures and tools were identical throughout the study, and the system was recalibrated as necessary to maintain accurate detection (approximately monthly). After calibration, Motive reported mean spatial triangulation errors of less than 0.05 mm throughout all experimental sessions. For kinematic tracking, marker detection thresholds were set to minimize spurious detection of non-marker objects (e.g., the mouse’s eye or snout). The threedimensional reach position data were saved in MATLAB (2015b) along with the time of reach success, defined as bringing the pellet into the behavioral arena. Reach success was monitored manually by the experimenter through a keypress function in MATLAB.

### Behavior

Animals were trained on a skilled reach task (Azim, Jiang, et al., 2014; Becker & Person, 2019; Whishaw, 1996). Briefly, animals were food restricted to 80%–90% their initial body weight and monitored daily for weight gain or loss as well as signs of distress. They were accommodated to the behavioral arena, consisting of a custom plexiglass box with a 0.9 cm opening, and subsequently trained to reach for 20 mg pellets of food (BioServ #F0163, San Diego, CA). Pellets were placed on a cylindrical pedestal located 1 cm away, 2 cm above the bottom of the behavioral arena and 0.5 cm left-of-center to encourage reaching with the right hand. The pellet pedestal (0.5 cm diameter) was designed to avoid physical interference with outreach or return limb reach trajectories and required the animals to skillfully grab the pellet to retrieve it. On training day 1, the pellet location was moved close enough to the arena opening to be retrieved with the tongue, and was slowly moved farther away on subsequent days to encourage reaching with the limb. All animals learned to accomplish the task using the right hand. Animals were considered ‘trained’ when they could successfully retrieve 30 pellets in a 20 min behavioral session (N = 26; range of 1 to 10 days of training from first reach).

For data collection, animals were lightly anesthetized with isofluorane for approximately 30 seconds in order to affix the tracking marker to the paw and allowed to recover for at least 5 minutes prior to beginning a behavioral session. Pellets were manually placed on the pedestal by the experimenter using a non-infrared-reflective pair of forceps to prevent detection by the tracking system. Sessions lasted until 30 pellets were successfully retrieved or 20 min were spent in the arena, whichever came first.

For head-fixed animals, mice were anesthetized with a ketamine/xylazine cocktail (100/10 mg/kg) and a bolus of lidocaine (2%) was injected subcutaneously under the scalp. A small incision over the scalp was made to clear and clean the surface of the skull, and a custom-made aluminum head plate centered on Bregma was affixed to the skull using luting (3M, St. Paul, MN) and dental acrylic (Teet’s cold cure; A-M Systems, Sequim, WA). Following head-plate affixation surgery, mice were allowed to recover a minimum of 5 days before beginning food restriction. When mice reached their target weight (80-90% of starting weight), they were habituated to head fixation. Mice were attached to head bars with their body and rear legs resting in a plastic tube and their front paws supported by horizontal bar (Heiney et al., 2014). Pellets were moved close enough so that they could be retrieved with the tongue. Once mice could consistently retrieve pellets, the pellet was moved progressively further away and positioned on the right side of the mouse to encourage reaches with the right hand (location 2.4 cm out, 1.5 cm up, and 0.1 cm lateral from origin). Mice were trained for 20 minutes per day until they achieved a 50% reach success rate (defined per pellet instead of per reach) 3 days in a row at which point they were considered fully trained (N = 4; range of 5 to 10 days of training from first reach).

### Data analysis

Data were analyzed with custom-written functions in MATLAB, as described previously (Becker and Person 2019). Time-stamped three-dimensional paw position data were cleaned, filtered, and segmented into discrete reach events. In the data cleaning stage, we corrected for spurious object detection (< 3% of data frames detected more than one marker) by conducting a nearest-neighbor analysis to differentiate the marker from other objects. Next, we filtered continuous kinematic data captured throughout each entire behavioral session, including while the animal freely moved in the behavioral arena and performed reaches. Since the marker could become hidden from view between reaches, we linearly interpolated missing samples, applied the filter (4th order, 8 Hz low-pass Butterworth) (Yu et al., 1999), and then removed interpolated data. Reaches were segmented from the continuous data by applying a positional threshold corresponding to marker positions just outside of the behavioral arena. From the points of threshold crossings out and back, we looked backward and forward in time to include marker data in which the paw continuously moved in the outward or inward direction, respectively, capturing the whole movement from inside the box to endpoint to return back into the arena. For most kinematic analyses, reaches were clipped at the first local maximum in the outward direction (i.e. reach endpoint) that occurred outside of the behavioral arena. Thus, unless otherwise stated, the return phase of the reach was excluded from numerical analysis.

Reach velocity and acceleration were calculated in each dimension as the numerical gradient of reach position and velocity data, respectively. Speed was the defined as the Euclidean magnitude of velocity values at each timepoint (2-norm). We defined the endpoint of the reach as the first local maximum in the outward direction. To calculate cumulative endpoint error, we took the sum of the standard deviation of reach endpoints across the three dimensions. Speed profiles were constructed by averaging time-interpolated velocity vectors at 10 ms intervals over a 400 ms window aligned to time of peak velocity. Asymmetry of the speed profile was calculated as the area under the first half of the speed profile divided by total area, with values of 0.5 representing perfect symmetry, values greater than 0.5 representing skew towards the first half of the reach, and values less than 0.5 representing skew towards the second half of the reach. Reach distance was calculated as the cumulative difference between adjacent reach data points in 3-dimensions (arc length). Duration was simply the difference in time between the first and last reach data points that resulted from the segmentation and first-endpoint clipping procedure described above. Mean deceleration was calculated by averaging outward acceleration values from the time point of peak velocity until the endpoint of the reach. For each of the above measures, we calculated within-animal averages across reaches, and then took the average across animals to report the population mean.

For trial-over-trial analysis, we started with unclipped reaches that included the full trajectory from outreach to return back inside the behavioral arena. We used the MATLAB ‘FindPeaks’ function to select reaches with more than one endpoint (‘attempts’), which we defined as peaks in the outward position vector that occurred outside of the behavioral arena and had prominence values (a measure of peak height) of at least 5 mm. Using this criterion, we analyzed the endpoints of first and second attempts relative to each other. We also used the same criteria to extract the final endpoint location of successful reaches, which we used as a proxy for accuracy, reasoning that the final endpoint represented the movement segment that resulted in successful grasping and retrieval of the pellet.

We used a random forest model to predict success of each reach based on kinematic variables (Breiman, 2001). We analyzed first-reversal clipped reaches (described above), and redefined successful reaches as reaches that succeeded on the animal’s first attempt. We limited our analysis to mice that had at least 10 first-attempt successful reaches, for a total of 3,239 trials across 17 mice. 70% of this data was used to train the model, and the remaining 30% was used to test predictions. Accuracy of the model was assessed for each animal independently, as well as for all reaches pooled together. To reduce the number of variables in the random forest model and have a consistent number of timepoints across reaches, we decomposed the reach for each kinematic variable (e.g. position out or velocity up) using a size K B-spline basis, where K was chosen to be 4. Using this approach, the kinematic variables for each reach can be represented using the 4 B-spline coefficients from this decomposition rather than all original timepoints. For example, for a reach with 95 original timepoints, 4 B-spline coefficient values are used to represent the outward position trajectory, while 4 different coefficient values represent the upward velocity trajectory. The 4 B-spline coefficients split the data into different time segments of the reach, where the first and fourth coefficients represent the first and last segments of the reach, respectively. These B-spline coefficients, as well as trial time length, were used as variables in the predictive model. Because there are 9 kinematic variables (upward, outward, and lateral directions each for position, velocity, and acceleration) and 4 B-spline positions for each kinematic variable, this amounts to 9 x 4 + 1 = 37 total variables in the model. Our approach has several advantages over using unprocessed trajectories, including better predictive capacity (Pfisterer et al., 2019), lower correlation between variables, and a reach-segment interpretation of the results. Random forest prediction modeling was performed in R using the package mlr (version 2.17.1, https://mlr.mlr-org.com/).

Minimum-jerk reach kinematics were generated using parameters from first-reversal clipped reaches (described above). The time and positions in each dimension (outward, upward, lateral) at the start and end of each clipped reach were used to compute minimum-jerk trajectories based on the previously described mathematical relationship:

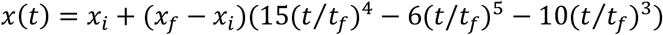

where *x*(*t*) is the position at time *t, x_f_* is the position at *t* = 0, and *x_f_* is the position at the final time point *t* = t_f_ (Flash & Hogan, 1985). Computing the derivative of the above equation yielded velocity in each direction. Speed was calculated from velocity magnitudes as described above.

Statistical tests were conducted in MATLAB using p < 0.05 criterion for significance. Parametric statistical tests were used only if data passed a Komolgorov-Smirnov test for normality. Otherwise, nonparametric tests were used. Paired tests were used for across-animal comparisons to account for interindividual differences in mean values. For most parameters, we report both within-animal metrics (e.g. correlations) and across-animal metrics (grand mean values). Simple linear regression was used to determine lines of best fit. Differences in within-animal variability were conducted using Levene’s test.

## Acknowledgements

We thank members of the Person lab for their insightful comments on the manuscript. Samantha Lewis and Elena Judd provided technical assistance. We thank Dr. Michael Hall and the Neuroscience Machine Shop for behavioral apparatus construction. This work was supported by National Research Service Award Individual Predoctoral fellowship (F31) NS103328 to M.I.B; F31 NS113395 to D.C; and grants R01 NS114430 and NSF Career 1749568 to A.L.P.

## Author Contributions

M.I.B. conceived the study, collected freely-behaving data, performed data analysis, made figures, and wrote the paper. D.C. collected head-fixed data, implemented and analyzed the minimum jerk model, and edited the paper. J.W. designed, implemented, and analyzed the random forest model, and edited the paper. A.L.P conceived the study and wrote the paper.

